# More than meets the eye: diversity and geographic patterns in sea cucumbers

**DOI:** 10.1101/014282

**Authors:** François Michonneau, Sarah McPherson, P. Mark O’Loughlin, Gustav Paulay

**Affiliations:** Florida Museum of Natural History, University of Florida, Gainesville, FL 32611-7800, USA; Marine Biology Section, Museum Victoria, GPO Box 666, Melbourne, Victoria 3001, Australia

## Abstract

Estimates for the number of species in the sea vary by orders of magnitude. Molecular taxonomy can greatly speed up screening for diversity and evaluating species boundaries, while gaining insights into the biology of the species. DNA barcoding with a region of cytochrome oxidase 1 (COI) is now widely used as a first pass for molecular evaluation of diversity, as it has good potential for identifying cryptic species and improving our understanding of marine biodiversity. We present the results of a large scale barcoding effort for holothuroids (sea cucumbers). We sequenced 3048 individuals from numerous localities spanning the diversity of habitats in which the group occurs, with a particular focus in the shallow tropics (Indo-Pacific and Caribbean) and the Antarctic region. The number of cryptic species is much higher than currently recognized. The vast majority of sister species have allopatric distributions, with species showing genetic differentiation between ocean basins, and some are even differentiated among archipelagos. However, many closely related and sympatric forms, that exhibit distinct color patterns and/or ecology, show little differentiation in, and cannotbe separatedby, COI sequence data. This pattern is much more common among echinoderms than among molluscs or arthropods, and suggests that echinoderms acquire reproductive isolation at a much faster pace than other marine phyla. Understanding the causes behind such patterns will refine our understanding of diversification and biodiversity in the sea.

## 1 Introduction

Estimates for the number of species on Earth are poorly constrained because sampling is challenging, and our understanding of species limits for most groups limited. Most metazoan species are delineated based on morphological differences, however, these morpho-taxonomic delineations have been challenged by the advent of molecular data that have revealed complexes of cryptic species ([1, 2, 3, 4]) and elucidated intra-specific polymorphism [5]. In particular, DNA barcoding emerged as a powerful technique to identify species, discover new species, and clarify species limits.

The importance of barcoding for species identification has been revived with the development of metabarcoding: the DNA-based community analysis of environmental samples. Second generation sequencing technologies allows the parallel sequencing of many gene copies, and can therefore be used to identify a diversity of species of small size (e.g., meiofauna in marine benthic sediments), lacking distinguishing morphological features (e.g., larvae in the plankton), or digested in the gut of their predators [6].

The success of species identification with these approaches relies on a comprehensive database to match sequences to be identified (unknowns) to reference sequences. Exact matching can only occur when the reference database comprehensively captures both the intra-specific genetic variation and the diversity of species [7]. In this case, the error rate will only depend on the proportion of species that share haplotypes. If the unknown sequence is not in the reference database, it could either represent an unsampled haplotype of a sampled species, or an unsampled species. Assigning these unmatched sequences to an unsampled species has typically relied on the “barcoding gap”, the potential absence of overlap between intra-specific and inter-specific distances [7]. If the unknown sequence diverged by a genetic distance above a threshold (typically 10 times the average intraspecific distance, or often set at 2-3%, see [1]), it would be attributed to a new species. However, intensive sampling typically revealed that intraspecific variation and interspecific divergence (based on accepted taxonomy of a group) overlap, i.e., there is no “barcoding gap” [8]. Additionally, differences in rates of molecular evolution among species make the application of a single threshold for species delineation challenging [7].

For species delineation, one of the greatest limitations of single locus approaches like barcoding, is that the gene tree (genealogy) may not reflect the species tree [9]. Including independent characters (e.g., nuclear loci, morphological, behavioral, ecological, geographical) in analyses provides a mechanism for testing for incongruences between gene trees and species trees. Reciprocal monophyly in two or more independent characters demonstrates lack of gene flow and delineates lineages with independent evolutionary histories. Such lineages are termed evolutionary significant units (ESU) [10], and can be treated as species-level units. However, they do not necessarily represent biological species, and their reproductive isolation can only be assessed when ESUs occur in sympatry.

When lineages identified with genetic data are associated with clear differences in morphology, morpho-taxonomic delineations are reinforced. Discordance between genetic data and morphological traits can occur when (1) morphological change has been insufficient to separate species, even though genetic data suggest isolation; (2) morphological change associated with speciation was so rapid that mitochondrial lineages have not fully sorted; (3) genetic divergence in the marker (e.g., mtDNA) has been erased by introgression; (4) morphological polymorphism does not reflect species limits (see [9, 11] for reviews). These situations might have undesirable effects on the accuracy of species identification, but they might provide insights into the biology of the species.

Many groups of marine organisms remain poorly explored or understood taxonomically [12], because the morphological characters traditionally used to delineate species frequently vary little among related species [13]. Molecular assessment of species limits in marine organisms has often led to recognition of high levels of cryptic diversity (e.g., [14, 15]), even for well-known species (e.g., [2]). These patterns have challenged the paradigm that the high dispersal potential of the larval stage in marine invertebrates limits their diversification [16].

The Indo-Pacific is the richest marine biogeographic region, and the origin of this diversity remains contentious. The paucity of obvious barriers to gene flow in the Indo-Pacific in combination with high levels of cryptic diversity, questions the importance of physical barriers to drive diversification in marine organisms. Yet, allopatric ranges predominate among young sister species (e.g., [14, 2, 17]). Exceptions are gaining attention, some pointing to a strong role for selection [18].

Here, we assembled one of the largest and most comprehensive (geographically and taxonomically) barcoding dataset. This dataset allows us to investigate levels of cryptic diversity and the geographic patterns of species diversity in sea cucumbers. Specifically, we ask: (1) How do estimates of species boundaries and diversity based on DNA sequence data compare with our current morpho-taxonomic assessment of sea cucumbers? (2) What is proportion of species that share haplotypes, or are not monophyletic, and how does it impact the identification success using DNA barcoding in this group? (3) How does the type of larval development influence the geographical distribution and genetic differentiation of species? (4) How does the frequency of sympatry change over time among sister species?

Sea cucumbers occur in most marine ecosystems, at all latitudes, from the intertidal to hadal depths. They constitute the largest invertebrate fishery in the tropical Pacific, and stocks are fully exploited to depleted throughout the tropics. Yet, species level taxonomy is poorly understood, with large, commercially important species still regularly described (e.g., [19]). In sea cucumbers, species-level taxonomy has almost entirely relied on the shape of ossicles, microscopic calcareous secretion permeating their tissues. Ossicles show substantial diversity and variability, whilst variation in the simple anatomy of these animals is limited at lower taxonomic levels. Ossicles cannot always be used to tease apart species, and intra-specific variation is poorly characterized since most descriptions illustrate ossicles from a single individual. Live coloration, which has not been traditionally used in species descriptions, is variable and can be indicative of species limits [4]. The combination of poor taxonomic understanding, and a pressing need to understand species limits, make a large-scale barcoding study of sea cucumbers especially important.

In sea cucumbers, as with other marine invertebrates, dispersal typically occurs during the larval stage. However, even if larvae can potentially travel long distances, their ability to maintain gene flow over long distances has been debated. If dispersal is spatially restricted, a pattern of isolation-by-distance (IBD), where genetic differentiation increases with geographical distance is expected. Larvae can be planktotrophic (they feed in the plankton), or lecitotrophic (they feed on yolk reserves). A few species of sea cucumbers are brooders, and release juveniles. The type of development and larvae influences the dispersal potential of the species, with brooders having little dispersal potential except in species prone to rafting, while species with planktotrophic larvae might disperse over longer distances. While the reproductive biology of most sea cucumber species has not been investigated, most species of Aspidochirotida and Apodida appear to have planktotrophic larvae, while all investigated species of Dendrochirotida have lecitotrophic larvae. Therefore, we hypothesize that (1) species with lecitotrophic development (Dendrochirotida) will show a steeper IBD pattern, and (2) smaller geographical ranges than species with planktotrophic development (Apodida and Aspidochirotida).

## 2 Materials and Methods

### 2.1 Sampling

Specimens were collected on snorkel, on SCUBA, or by dredging. Most were photographed while alive *in situ* or in the lab anesthetized in a 1:1 solution of sea water and 7.5% solution of magnesium chloride hexahydrate, then preserved in 75% ethanol. When possible tentacles were clipped, immediately put in 95-99% ethanol, and later used for DNA extractions. Specimens were deposited in the Invertebrate Zoology collections of the Florida Museum of Natural History, University of Florida (UF), Gainesville, FL, USA, while tissue samples are stored in the Genetic Resources Repository at UF.

Additional samples were obtained through collaborators either from newly collected material or from preserved specimens deposited at other institutions.

3048 individuals were identified and sequenced. They represent the worldwide diversity of sea cucumbers, with a particular focus on species of the Indo-Pacific and the Antarctic regions (Fig.1). The northern hemisphere and deep-water species outside of the Antarctic region are under-represented.

**Figure 1:**
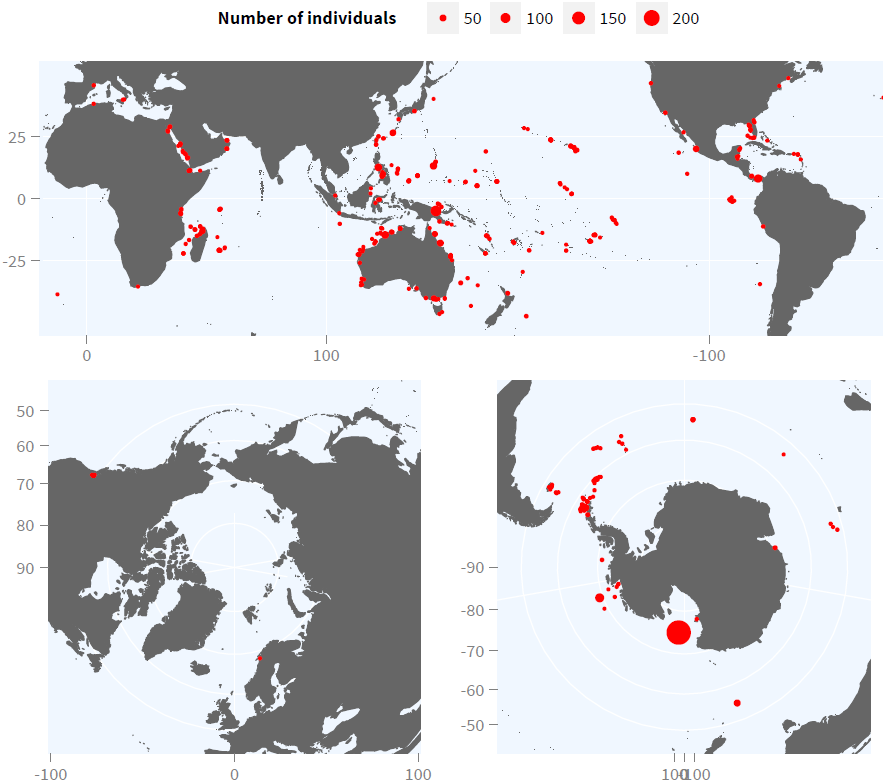
Locations sampled

### 2.2 Specimen identification

Most specimens were individually identified by GP, FM, Mark O’Loughlin and other taxonomic specialists. For most groups, extensive survey of primary literature was used to assign the most accurate names.

### 2.3 Sequencing and analysis

DNA was extracted according to one of the following three protocols: (1) DNAzol followed by QIAGEN PCR clean up kit; (2) organic extractions at the Smithsonian Institution using robotic facilities; (3) Omega EZNA Mollusc kit. In each case, the manufacturer’s protocol was used.

We amplified the 5’ end (655 base pairs) of the mitochondrial gene COI using the echinoderm primers developed by [20]. The sequencing was performed at the Interdisciplinary Center Biotechnology Research at the University of Florida, or at the Smithsonian Institution. Chromatograms were assembled and edited using Geneious 5.5.8 [21].

We kept all sequences that were more than 500 bp, and that did not include any stop or nonsense codons that may have resulted from incorrect base calling during the sequencing or the editing process. Sequences were aligned using MAFFT [22].

We generated Neighbor-Joining (NJ) trees from pairwise uncorrected distances as well as using the Kimura 2-parameter correction [23]. We assessed branch support using 200 bootstrap replicates. Additionally, we constructed a maximum-likelihood tree using RAxML 8.0.1 [24] with each codon position as a different partition, a GTR model of molecular evolution with a Gamma distributed rates of heterogeneity, and 500 bootstrap replicates.

### 2.4 Diversity

The size of the dataset restricts the methods that can be used to assess diversity, as computationally intensive methods such as the Generalized Mixed Yule Coalescent method [25] cannot be used here. Other programs cannot accommodate such large dataset, and failed to complete the analysis (e.g. PTP [26], AGBD [27]). We assessed species boundaries and diversity in three ways.

First, we used traditional morphological characters and current species concepts across the class to identify species to currently valid species names. For this assessment we used these names at face value, although we are aware of cryptic morphological variation in many species we have studied in depth, and such variation likely also exists in many less-studied forms. Overall, 89 morphospecies could not be assigned to a name. Some of the material included in this study was provided to us as tissues, and the specimens were not evaluated to be identified with confidence. For some groups, taxonomic understanding is limited, and species names cannot be assigned accurately. Therefore, a proportion of the morphospecies for which we could not assign a name might match already described species, but others are undescribed.

Second we used the evolutionary significant unit (ESU) concept (*sensu* [10]) to evaluate diversity within the holothuriids (= family Holothuriidae vs. holothuroids = class Holothuroidea). They were the most densely sampled and comprise almost half (47%) of our dataset; they are also the focus of our taxonomic work and are the best known to us at the morpho-species level to assign all individuals to ESUs. Individuals were assigned to ESUs based on differentiation in at least two independent characters (e.g., reciprocal monophyly in COI, morphology, ecology, restricted geographical range).

Third, we used mtDNA sequence data on its own for lineage delineation (mtLineages) based on thresholds [7, 1]. We grouped individuals that differed from each other by a genetic distance below a given threshold (“pairwise” method). This method is strictly based on genetic distance and does not take into account phylogenetic information. We also identified monophyletic groups from Neighbor-Joining trees, reconstructed from pairwise genetic distances (“clustering” method). Groups were defined based on coalescence depths that were below a threshold set *a priori.*

For each of these two threshold-based approaches, we used both uncorrected distances (*p*) and the Kimura two-parameter (K2P) model [23] to estimate genetic distances. The K2P model is very widely used when analyzing barcoding data, but a recent study indicated that *p* distances are probably more appropriate when comparing genetic distances between closely related species and may lead to higher rates of identification success [28]. We estimated the number of species recovered from our sample while varying the pairwise threshold value: 0.01,0.015,0.02,0.025,0.03,0.035,0.04,0.045,0.05,0.06, 0.07, 0.08. Additionally, we estimated the proportion of singletons (proportion of the number species represented by a single individual) for each of these threshold values.

To assess the accuracy of the pairwise and clustering methods, and the most appropriate threshold, we compared the manually delineated ESUs to the mtLineages estimated by these methods in the Holothuriidae. We tested which method (pairwise or clustering) and which threshold recovered most closely the manually delineated ESUs (closest estimated number and lowest error rate). We measured the error rate of the threshold-based approaches by recording the number of ESUs that were oversplit (one ESU included more than one mtLineage) or lumped (one mtLineage included more than one ESU) for each threshold.

### 2.5 Geography, diversity and diversification

#### 2.5.1 Range size and isolation-by-distance

To test whether mode of development had an impact on range size, we compared the maximum geographical distance between sampled individuals for each mtLineage as delineated with the threshold approach (4.5% threshold with clustering method, justification below).

To test for evidence of IBD we investigated the relationship between the maximum genetic distances within mtLineages and the maximum geographical distances sampled for each mtLineage represented by three or more individuals. If dispersal is spatially restricted, we expect genetic differentiation to increase with geographical distance. To assess if differences in the reproductive modes among the orders had an effect of the IBD pattern, we performed an analysis of covariance (ANCOVA) on the maximum genetic distance for each species as the response variable, the maximum geographic distance as the covariate, and the taxonomic order as the independent variable. We performed the analysis on the three most represented orders (Apodida, Aspidochirotida, Dendrochirotida), which are also the best understood in terms of larval development (Apodida and Aspidochirotida predominantly with planktotrophic larvae, Dendrochirotida with lecithotrophic larvae). The variances among the orders were homogeneous (Levene’s test, *F*(2, 257) = 1.272, *P* > 0.28). To assess if using the threshold-based approach might have influenced the results, we compared the model obtained with the model estimated from the manually delineated ESUs.

#### 2.5.2 Geography of diversification

If speciation occurs in allopatry, recently diverged species may retain allopatric distributions for a period of time. To investigate the geographic mode of diversification, we evaluated whether the geographical ranges of sister ESUs overlapped. We identified whether the ESUs were sister based on the maximum-likelihood tree. If the distributions of the ESUs could be inferred with a polygon (i.e., consisted of at least 3 occurrences), we classified their distributions as: “parapatric” if less than 10% of the range of the ESU with the smallest range overlapped with the ESU with the larger range, “sympatric” if their ranges overlapped by more than 10%, and “allopatric” otherwise. If only one ESU had enough data to estimate its distribution with a polygon, and points were available for its sister ESU, we considered their distribution allopatric if none of the points were included in the polygon, and sympatric otherwise.

To investigate the relative contributions of geographic barriers to the formations of ESUs within the Indo-Pacific, we recorded the approximate location of the boundaries separating allopatric sister ESUs.

## 3 Results

### 3.1 Diversity

Among the 3048 individuals sequenced, 389 identified species were included. All orders of sea cucumbers were represented. The Aspidochirotida, and in particular the Holothuriidae were the most comprehensively sampled (47% and 33% of the total number of individuals respectively), with all genera and subgenera of the Holothuriidae represented, and 65% for the currently accepted species.

Manual delineation of the ESUs in the Holothuriidae revealed 194 ESUs, 49% more than named species (N=130). About one quarter (N=31) of named species were complexes, each including between 2 and 13 ESUs. Additionally, 24 ESUs appeared to be undescribed species (not directly referable to a named species). 24% of the ESUs were singletons. Most but not all named species were reciprocally monophyletic (6.1%, N=9 of the ESUs were non-monophyletic).

The distance to the nearest neighboring ESU was greater than the maximum intra-ESU distance for 91% (N=136) of the manually delineated ESUs (Fig. 2). The minimum inter-ESU distance varied greatly, but was > 2% for 91% (N=136).

**Figure 2:**
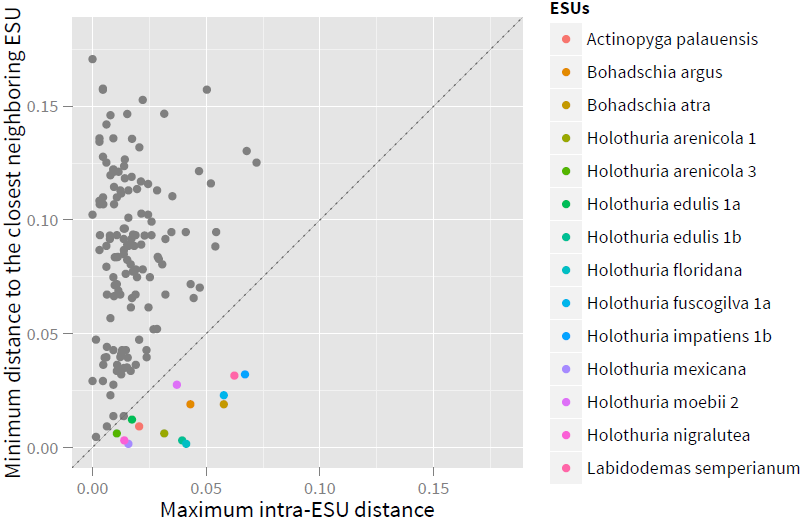
Comparison of interand intra-ESU uncorrected COI distances. Foreach ESU we compared the maximum intra-ESU genetic distance with the minimum inter-ESU distances, ESU above (gray) the 1:1 dotted line can be easily delineated with the barcoding gap, the ESUs falling below the line (colored) cannot.

On the entire dataset, regardless of the method or the distance threshold used, the number of estimated mtLineages was greater than the number of morpho-species. The pairwise distance method estimated lower numbers of mtLineages at the same thresholds than the clustering method, and was less sensitive to the type of distance used (Fig. 9, 10).

The threshold approaches estimated similar numbers of mtLineages and proportions of singletons as for the ESUs with thresholds between 3% and 5% depending on the approach and the type of distance used (Fig. 9).

The greatest correspondence among threshold methods compared to ESUs delineated was with the clustering method and a threshold of 4.5%; this method delineated 19% (N=37) of the ESUs incorrectly. The number of mtLineages estimated with this method was slightly lower than the number of ESUs (183 and 194 respectively) as more ESUs were lumped than split (23 and 14 respectively). Thus, we chose the clustering method, using uncorrected distances and a 4.5% threshold to assess mtLineage diversity in subsequent analyses (Table 1).

**Table 1.** Number of named morpho-species sampled (# sampled), number of accepted species (# accepted), and number of mtLineages (# mtLineages) estimated with the clustering method and a 4.5% threshold for each family and each order of sea cucumbers. There were 194 ESUs delineated for the Holothuriidae. Not all families were sampled, thus totals in some orders are more than sum of family diversities. This classification does not include modifications proposed by Smirnov [29]. Estimations of the number of mtLineages is based on different datasets for families and orders, thus totals for some orders may differ because of taxonomic uncertainty (samples identified at the order level but not at the family level), or differences in lineages delineation when the entire order is considered

| Order | Family | # accepted | # sampled | # mtLineages |
| --- | --- | --- | --- | --- |
| **Apodida** |  | **307** | **34** | **64** |
|  | Chiridotidae | 81 | 14 | 26 |
|  | Synaptidae | 173 | 20 | 37 |
| **Aspidochirotida** |  | **386** | **186** | **275** |
|  | Holothuriidae | 200 | 130 | 188 |
|  | Mesothuriidae | 33 | 9 | 14 |
|  | Stichopodidae | 32 | 26 | 29 |
|  | Synallactidae | 121 | 21 | 43 |
| **Dactylochirotida** |  | **42** | **2** | **6** |
|  | Ypsilothuriidae | 15 | 2 | 5 |
| **Dendrochirotida** |  | **710** | **127** | **213** |
|  | Cucumariidae | 294 | 62 | 91 |
|  | Paracucumidae | 3 | 4 | 4 |
|  | Phyllophoridae | 188 | 20 | 50 |
|  | Psolidae | 132 | 23 | 39 |
|  | Sclerodactylidae | 87 | 18 | 26 |
| **Elasipodida** |  | **158** | **27** | **36** |
|  | Deimatidae | 14 | 2 | 5 |
|  | Elpidiidae | 94 | 11 | 13 |
|  | Laetmogonidae | 23 | 8 | 11 |
|  | Pelagothuriidae | 2 | 1 | 3 |
|  | Psychropotidae | 25 | 5 | 5 |
| **Molpadida** |  | **101** | **13** | **23** |
|  | Caudinidae | 34 | 3 | 7 |
|  | Molpadiidae | 63 | 10 | 17 |
| **Total** |  | **1704** | **389** | **614** |

### 3.2 Geography, diversity and diversification

#### 3.2.1 Range size and isolation-by-distance

Apodida and Aspidochirotida had larger range sizes than the Dendrochirotida (Fig. 4, median of the range sizes of 5242 km, 4199 km, 710 km respectively).

**Figure 3:**
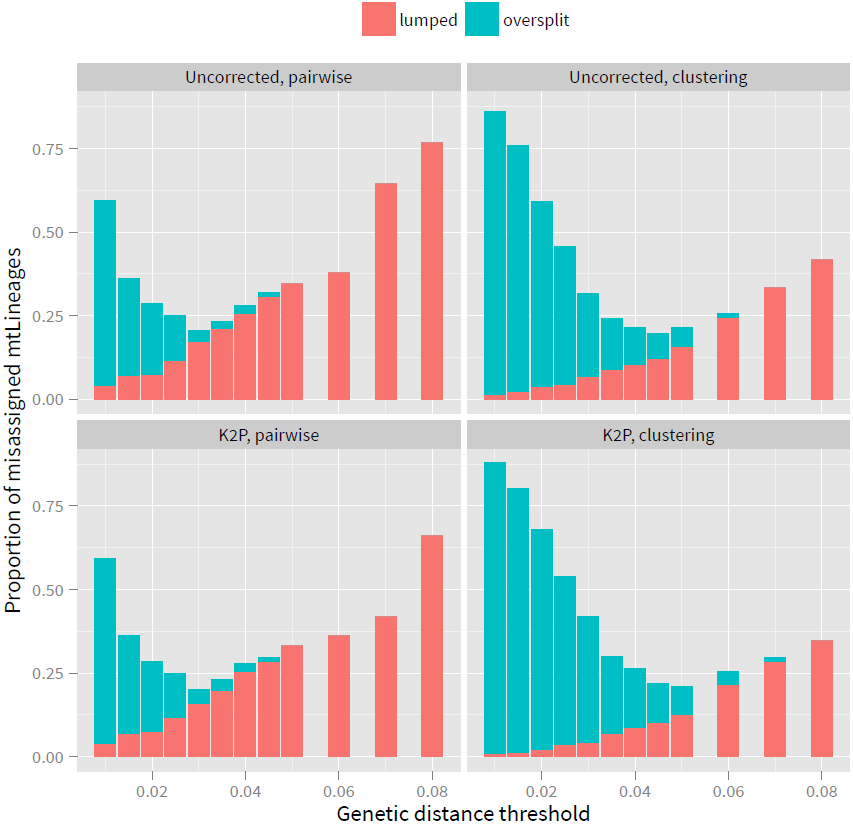
Proportion of mtLineages incorrectly assigned compared to ESUs delineated on the Holothuriidae using pairwise and clustering methods, based on uncorrected and Kimura 2parameter genetic distances (K2P). mtLineages can be incorrectly assigned because of oversplitting (one ESU represented in two or more mtLineages), or lumping (one mtLineage is comprised of several ESUs). The lowest error rate is with the clustering method based on raw distances and a threshold of 4.5%.

**Figure 4:**
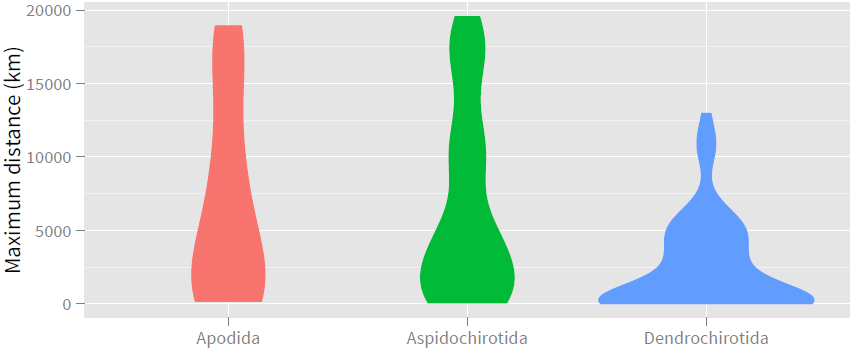
Violin plot (widths represent the density estimate) of the maximum range sizes for the Apodida, Aspidochirotida, Dendrochirotida. Note that most Dendrichirotida have smaller distributions.

We recovered a signal for isolation-by-distance (IBD) in each of the three orders included in the analysis (ANCOVA, *F*(1, 258) = 45.1, *P* < 0.001, Fig. 5, Table 2). The slopes and the intercepts were similar across the three orders (slope interaction: *F*(2,254) = 2.05, *P* = 0.13; intercept interaction: *F*(2,254) = 0.949, *P* = 0.39), and were not significantly different from the coefficients estimated on the ESUs (intercept: 0.012, SE=0.0012, slope: 8.4e-07, SE=1.425e-07).

**Figure 5:**
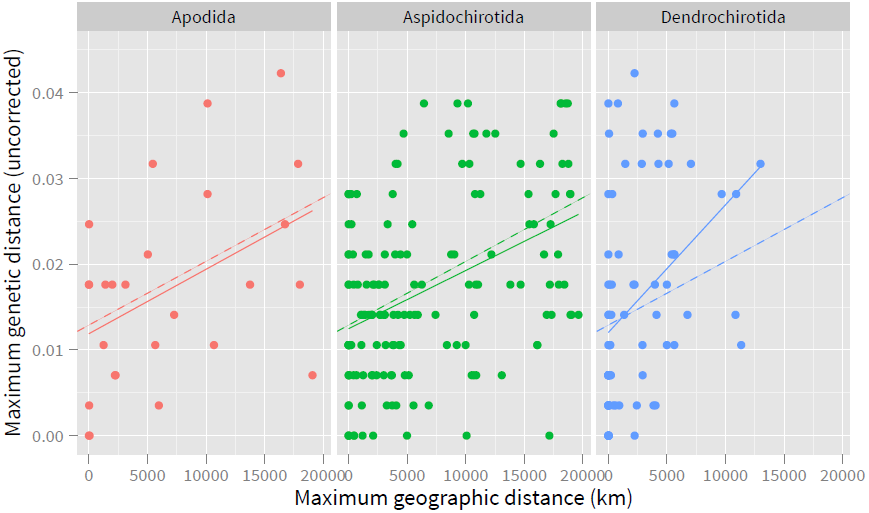
Maximum genetic distance and maximum geographic distance for mtLineages identified with the clustering approach 4.5% threshold represented by at least 3 individuals. Dashed lines represent the model estimated by an ANCOVA using the order as the independent variable. Solid lines represent the linear model for each order.

**Table 2.** Coefficients of the regression between maximum genetic distances and maximum geographic distances for all ESUs identified with the clustering method with a threshold of 4%, represented by 3 or more individuals. See Fig. 5.

|  | Estimate | Std. Error | t value | Pr(>\|t\|) |
| --- | --- | --- | --- | --- |
| (Intercept) | 0.01289 | 0.0008891 | 15 | 2.966e-35 |
| maxGeoDist | 7.426e-07 | 1.105e-07 | 6.7 | 1.169e-10 |

#### 3.2.2 Geography of diversification

We compared the geographic ranges of 40 pairs of sister ESUs within the Holothuriidae. Most had allopatric distributions (82%), the remaining were sympatric (15%), or parapatric (2.5%).

There was no evidence that sister ESUs occurring in sympatry were older than ESUs occurring in allopatry, as half (N=3) of the sympatric sister ESUs showed genetic distances lower than the median of the distances observed for allopatric ESUs (Fig. 6).

**Figure 6:**
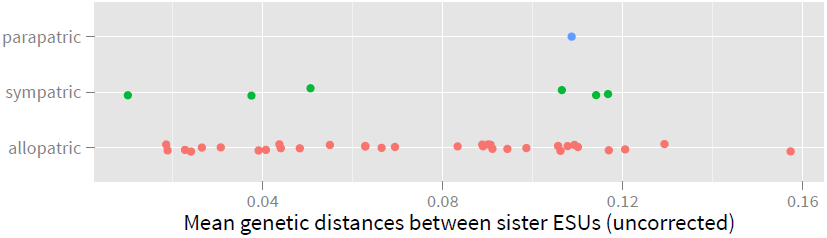
Mean genetic distances between sister ESUs occuringin sympatry, allopatry and parapatry.

Sister species with allopatric ranges tended to straddle well-known biogeographic boundaries. Thus 42% of the allopatric pairs separated between the Indian and Pacific Oceans, 18% between the Red Sea/ Arabic peninsula and the rest of the Indian Ocean, and 12% between the Hawaiian archipelago and other Pacific islands. Other barriers separated endemics restricted to remote locations and/or in areas with distinct ecological conditions (Galapagos, French Polynesia, Eastern Pacific, Oman).

Genetic divergences across the main biogeographic boundaries were variable, and suggestive that isolation was asynchronous (Fig. 7). For the three barriers investigated, genetic distances seemed to form two groups separated by at least 2.5% divergence.

**Figure 7:**
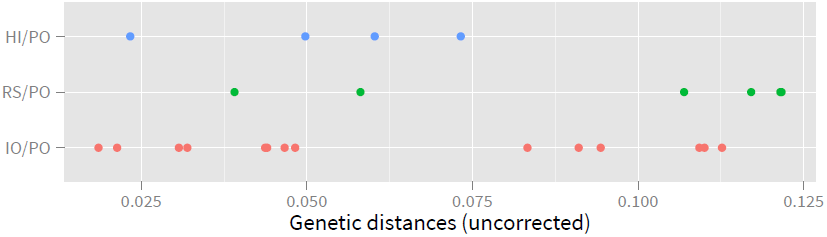
Mean genetic distances between sister ESUs separated by the Indian Ocean (10) and the Pacific Ocean (PO), the Red Sea (RS) and the Pacific Ocean, and Hawaii (HI) and the Pacific Ocean

## 4 Discussion

### 4.1 Diversity

Sea cucumber diversity is underestimated because many species currently delineated are complexes of several ESUs and because undescribed species are common. Regardless of the approach used to estimate diversity from genetic data, we recovered more species than we identified based on accepted taxonomic names. In the best data (i.e., the manual assessment of the ESUs for the Holothuriidae), the diversity is underestimated by at least 50%, a proportion that will increase with further sampling as evidenced by the high proportion of singletons. This family is the best studied: it is easily accessible as it inhabits shallow waters, species are large, many are economically important, and most species have widespread geographical distributions. Other families likely have higher levels of unrecognized diversity.

Such levels of unrecognized diversity were not expected, and contrast with other marine invertebrates that have received more taxonomic attention (8 out of 263 sampled ESUs corresponded to previously unrecognized species or sub-species in cowries [7], and 9 out of 52 ESUs for hermit crab [17]). Sea cucumbers are generally considered to be well understood taxonomically, as variation in taxonomic characters appear to be limited and documented. However, our data show that many of the current morphospecies concepts, even in the best known family (i.e., Holothuriidae), are in many cases inaccurate and do not reflect the species limits suggested by genetic data. Our studies (e.g., [30,4]) and others (e.g., [31]) have emphasized that genetic data can help redefine the morphological traits that can be used to delineate species. Notably, differences in color patterns emerge as useful indicators of species limits. A taxonomic overhaul, guided by genetic data, is needed to clarify species limits in all sea cucumbers.

### 4.2 Species limits

The ESU approach provides guidelines for identitying of biological species. Even if correlation between independent traits does not guarantee that ESUs are biological species, the concept demonstrates lack of gene flow and provides a powerful indirect test for reproductive isolation [7]. More than 94% (N=138) of ESUs identified for the Holothuriidae are reciprocally monophyletic in COI and are morphologically distinct, making them natural targets for taxonomic revision. The ESUs that are not reciprocally monophyletic for COI sequences are either (1) monophyletic but their closest neighbor is separated by distances lower than the maximum intra-ESU distances (10 out of 15 ESUs), or (2) not monophyletic. Interestingly the 5 ESUs that are not monophyletic all belong to the subgenus *Halodeima:* 3 ESUs of *Holothuria edulis, H. mexicana* and *H. floridana.* Additionally, 2 species of *Actimpyga,* each represented by a single individual are paraphyletic with *Actinopyga* species (*A.flammea* and *A. caerulea* are indistinguishable from *A. palauensis).* Other species outside the Holothuriidae are also found to not form reciprocally monophyletic lineages (e.g., the *Stichopus “variegatus”* complex in the Stichopodidae).

Funk and Ormland [9] estimated that 23% of the 2,319 animal species they surveyed were nonmonophyletic in their mitochondrial genealogies. Such species may not be distinguishable with DNA barcoding, and this high proportion has been used as an argument against the method (e.g., [32]). Funk and Omland found that the proportion of polyphyletic species was negatively correlated with the intensity of study, indicating that inadequate taxonomy was an important source of this error, and may have inflated their results. Here, we recovered similar rates of non-monophyly when considering currently defined morphospecies (17%, N=90), but much lower rates when using ESUs defined by integrative taxonomic study ESUs (6.1%, N=9). Careful taxonomic investigation is thus required to ensure accurate estimation of non-monophyly, and to highlight biologically interesting cases.

Three hypotheses can explain the absence of reciprocal monophyly in these well-studied morphologically distinct species. First, these species might have diverged very recently leading to incomplete lineage sorting. During the sorting of gene copies that split before the divergence of the species, the lineages will first appear polyphyletic, then paraphyletic, and will finally form reciprocally monophyletic groups [33]. The time taken to pass through these stages will depend on how rapidly drift will remove the apparently incorrectly sorted lineages. Second, species could have acquired mitochondrial copies from each other through introgressive hybridization, masking earlier differentiation. Third, the putative morphospecies could represent phenotypic variation within a single species.

Introgression appear uncommon among marine invertebrates (see [14] for examples of introgression in *Australium),* but has been reported in echinoderms (e.g. [34, 35]). Two instances of hybridization between sea cucumber species have been documented [36,31]. However, in both cases hybrids do not appear to backcross with their parental species limiting the possibility for genomic introgression. Teasing out the processes that are leading to non-monophyletic patterns observed in COI requires multilocus dataset, which analyzed with the multicoalescent [37] might provide support for reproductive isolation among the putative species (see [4]). Recently developed methods that use single nucleotide polymorphisms (SNP) across the genome [38] appear as powerful tools to explore species limits among closely related taxa. These approaches remain expensive and time-consuming but come in complement of single-locus studies to untangle the lineages that are not monophyletic.

### 4.3 Methodological considerations

Modern taxonomy seeks to align taxa with evolutionary groups, however, while evolutionary groups can overlap, interbreed, be nested within one another, taxa are created to reflect distinctiveness (most commonly in morphology and/or evolutionary history depending on the species concept considered) [39]. This difference in the nature of boundaries between taxa and evolutionary groups creates conflict during the delineation process. In particular, attempting to delineate reproductively isolated taxa on the basis of genetic differentiation of a single marker will always be associated with some rate of failure if two or more species share haplotypes. The goal is therefore to find a method that captures best the range of genetic differentiation observed across species while identifying conflict in the data pointing to species that will be problematic to delineate, and might provide interesting biological insights.

Beyond species that share haplotypes, analytical challenges with methods estimating species limits only from genetic datasets, is to account for variation in both intraspecific and interspecific (to the closest species) genetic distances. Higher than average intraspecific distances will lead to oversplitting by recognizing divergent haplotypes as species, while lower than overage distances to the nearest neighbor will lead to lumping. In our study, the lowest error rate associated with threshold-based delineation was similar across the methods used (about 20%), but the pairwise approach was more conservative by estimating higher lumping rates and fewer lineages than the clustering method. By only delineating lineages based on distances and no phylogenetic information, the pairwise approach lumped reciprocally monophyletic lineages separated by low inter-ESUs genetic distances.

Methods that attempt to account explicitly for variation in intra- and inter-specific distances, such as GMYC [25, 40], or ABGD [41], might provide more accurate estimates. However, the available softwares that implement these approaches are not designed to analyze dataset as large as in this study. GMYC is also sensitive to the method used to obtain the ultrametric tree needed for the analysis ([4]), which could have large effects on analyses of dataset like ours. Few studies have compared the performance of GMYC in groups well-understood taxonomically, but Talavera *et al.* [42] found that it led to a 20% error rate in butterflies, which is comparable to what we found here using threshold-based approaches.

### 4.4 DNA barcoding for species identification in sea cucumbers

One of the main applications of DNA barcoding is to facilitate species identification, especially when only non-diagnostic parts are available (e.g., when processed for trade [43], in larvae [44], for gut contents [45]). Accurate identification requires a comprehensive database of DNA sequences to capture the diversity of species, as well as intra-specific variation. If the sequences for the individuals to be identified have exact matches in the reference database, the accuracy of the identification will only depend on the proportion of species that share haplotypes. However, commonly, these unknowns will not be in the database, and the accuracy of a positive identification will also depend on the accuracy with which the sequence can be predicted to belong to a species included in the database, and the probability that species included in the database share unsampled haplotypes. The accuracy with which a sequence can be predicted to belong to a species included in the database is related to the proportion of species that are not monophyletic.

In our ESU dataset, if the unknown sequences have an exact match in our database, individuals can be identified with high accuracy through DNA barcoding, as only 7 out of 194 ESUs (3.6%) share haplotypes. If the unknown sequences belong to ESUs that are not reciprocally monophyletic, or that are characterized by higher intra-ESU distances than the distance to the closest neighboring ESU, they will likely be misidentified. Together with the non-monophyletic ESUs, they represent 8.2% (N=16 ESUs) of our ESU dataset.

Sequences without a match in reference databases are typically assigned to a sampled species if the sequences differ by less than a threshold, or are considered as unsampled species if the genetic distances are above this threshold [7]. This is typically done either in the context of pairwise genetic distances or in the context of the closest match in a Neighbor-Joining tree. Thresholds are estimated based on the extent of overlap between intra-and interspecific genetic variation among the species considered, the barcoding gap. One of the issues associated with this concept is that increased sampling reduce, and often eliminate, the barcoding gap by capturing closely related yet good species, or rare, divergent haplotypes that increase the overlap between intra-and interspecific variation [7, 32]. More recent methodological developments account for the distance to the nearest neighbor to determine whether a sequence represents an unsampled species [46], or use a machine learning approach to detect features in the sequence that allow species identification [47].

### 4.5 Geography of diversification

Understanding the geographic context of speciation can inform the nature and strength of evolutionary forces that are driving reproductive isolation [48]. Additionally, understanding the factors that limit how soon after diverging in allopatry species can co-occur, or how often – if ever species – might be able to diverge in sympatry, may contribute to the heterogeneity in the spatial distribution of species diversity. In oceans where organisms might disperse over large distances and barriers to dispersal are rare, opportunities for geographic isolation seem limited. However, species with shortlived larvae are expected to have smaller geographic ranges, and higher levels of genetic differentiation across their ranges.

We detected a consistent pattern of isolation-by-distance for the three orders we sampled more thoroughly despite differences in dispersal potential. Our results suggest that on average, ESUs sampled across larger geographical ranges show higher levels of genetic differentiation. Nevertheless some species show remarkable powers of dispersal, sharing haplotypes across the entire Indo-Pacific, from Africa to America. The type of larva does not seem to affect the amount of genetic differentiation expected across the range as evidenced by the common slope and intercept estimated across orders. However, the Dendrochirotida with their lecithotrophic larvae, have smaller range sizes than the orders with planktotrophic larvae. Both Apodida and Aspidochirotida have ESUs that span the entire Indo-Pacific while only a few Dendrochirotida have ESUs that span either the Indian or the Pacific oceans. These results suggest that the type of larvae influence patterns of diversity and modes of diversification.

Dendrochirotida species tend to have restricted ranges, with many species known only from single localities, and species within a genus are often strictly allopatric. In the Indo-Pacific, most species of Dendrochirotida are restricted to continental areas, with *Afrocucumis africana* being the only species that reaches the Central Pacific. This pattern could be explained by the combination of limited dispersal abilities that make the insular Pacific difficult to reach, and the suspension feeding mode of this group that cannot inhabit the oligotrophic environments that characterize oceanic islands [49].

However, as some Dendrochirotida species are panmictic across continental areas of the Indian Ocean, dispersal limitation seem unlikely lending support to the importance of ecology to explain the distribution of this group.

As species of Apodida and Aspidochirotida can maintain connectivity across very large distances, opportunities for allopatric speciation are limited. Nevertheless, most sister ESUs were found in allopatry indicating the importance of geographical isolation in the diversification of sea cucumbers. Allopatric speciation appears to be the most common mode in animals [48] and is also the most prevalent mode in marine organisms (e.g., [14, 17]). Well known biogeographic barriers have served to drive speciation as evidenced by allopatric sister ESUs in the Holothuriidae straddling these. Basinal separation between the Indian and the Pacific Oceans, and isolation of Red Sea/Arabia and Hawaiian archipelago were the three most important barriers for cladogenesis (Fig. 8).

**Figure 8:**
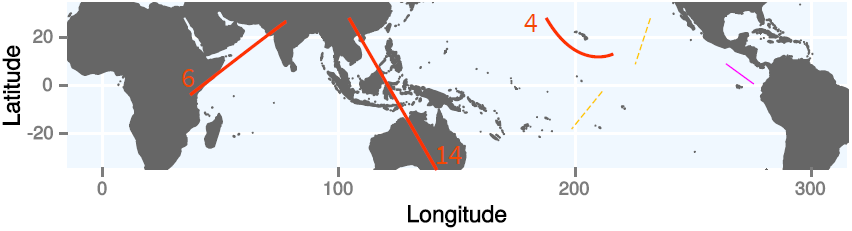
Approximate location of the boundaries between allopatric ESUs. Numbers indicate diversification events across these boundaries. Pink line 3 events, dashed yellow line 1 event.

The connection between the Indian Ocean and the Pacific Ocean was restricted several times throughout the Pleistocene [50] which could have contributed to the isolation and divergence of the species across the Indo-Pacific Barrier (IBP). The IBP is a recognized marine biogeographic barrier, and in a recent review, 15 out of 18 species of fish and invertebrates surveyed showed genetic differentiation across this barrier [51].

The Hawaiian archipelago and the Red Sea are well recognized areas of endemism in the IndoPacific [52, 17, 53, 54]. The Hawaiian archipelago is remote and its endemics are thought to have originated through peripatric speciation events [17, 54]. The Red Sea is separated from the Indian Ocean by a narrow strait, which in combination with areas of upwelling along the Arabian coast, likely limit population connectivity with the rest of the Indo-Pacific. This, in combination with its peculiar oceanographic conditions, maintains the distinctiveness of the fauna. However, the Red Sea experienced several salinity crises, the latest as early as *ca.* 19,000 years ago [53]. Therefore, the Red Sea fauna must have colonized the area from outside refugia following the salinity crises.

Understanding how these barriers drive speciation events across these biogeographic barriers is key to uncovering the mechanisms generating diversity within the Indo-Pacific. Three models have been proposed to explain how these barriers might be acting [55]. A model of classic vicariance emphasizes the importance of large-scale events (e.g., geological) that split an ancestral population into populations that diverge in isolation. A modified version of this model (“soft vicariance”) proposes that long-distance gene flow connects diverging populations that eventually become separated by changes in oceanographic conditions. Under a peripatric model, a new population is founded by the colonization of a small number of individuals from a larger source population [55]. Additionally, determining whether sister species separated by the same barrier, diverged concomitantly could inform on the relative importance of extrinsic factors (e.g., geology, climate) *vs.* intrinsic factors (e.g., selection) in generating diversity. Our analyses limit the inference on the nature of these barriers and whether they affected all species synchronously, however, the spread of genetic distances (Fig. 7) suggests multiple events. Determining the model of speciation that best explains the patterns of divergence would require additional analyses such as [56, 55]. Hodge *et al* [54] showed that coral reef fish species endemic to the Red Sea diverged steadily through time, while species endemic to the Hawaiian archipelago diverged in two pulses (0-3 million years ago [Mya] and 9-13 Mya). Our data for Hawaii is limited, but consistent with these results as the lowest divergence is for 2 species of the *Holothuria impatiens* complex with divergences estimated at 2 Mya ([4]).

Compared to studies on other taxonomically well-understood marine organisms, we recovered a higher proportion of sister ESUs occurring in sympatry for the Holothuriidae. For instance, sister ESUs are almost always allopatric in oceanic settings for molluscs (e.g., [14, 57, 58]) and arthropods (one out of 40 ESUs for hermit crabs in [17]). In contrast, we recovered several closely related and sympatric ESUs that exhibit distinct color patterns and/or ecology, show low-levels of differentiation in, or in some cases cannot be separated by, COI sequences alone. If these forms represent distinct species (as their morphology and biology suggest, a pattern we also found in other holothuroids and echinoderms), it would indicate that they have diversified more rapidly than most marine invertebrates. Additionally, in our study many closely related ESUs occur in sympatry, indicating that reproductive isolation can be completed rapidly. For instance, we showed how 3 ESUs in the *Holothuria impatiens* complex, occur in sympatry even though they diverged less than 2 millions years ago [4]. Four out of the eight complexes with three or more ESUs, have ESUs with overlapping ranges. This contrasts with other marine organisms, for which secondary sympatry in oceanic settings occurs slowly and takes at minimum 2 millions years but often more than 10 millions years [14, 59, 17, 60].

This difference could be the result of sympatric speciation, more rapid secondary sympatry (speciation occurred in allopatry, but the resulting species extended their ranges, and are now sympatric), or a sampling artifact (the actual sister species have not been sampled). If speciation in complete sympatry remains controversial, recent studies have shown that selection along depth gradients [61], host shifts [62] or other forms of selection (reviewed in [18]) might promote divergence even with some levels of gene flow.

Rapid acquisition of reproductive isolation could arise from premating isolation barriers such as differences in the timing of gamete release or gametic incompatibilities caused by the gamete recognition proteins (GRPs). GRPs are known to evolve rapidly and play a critical role in fertilization success of many free spawning marine invertebrates such as sea urchins [63, 64] and gastropods [65]. For instance, strong positive selection has been detected in the GRPs between closely related species of sea urchins in sympatry but not in allopatry [66], suggesting the role of these proteins in maintaining species limits. GRPs have not yet been identified in sea cucumbers but bindin is known from sea urchins [64] and sea stars [67] which indicates that it is plesiomorphic in the clade that sea cucumbers emerged from.

## 5 Conclusion

Our study shows that species diversity in sea cucumbers is underestimated by at least 50%. In many cases, DNA barcoding data appears useful to guide taxonomic decisions, as most unrecognized species differ in live coloration, but are currently not recognized as they do not vary in the taxonomic characters typically used to tease apart species.

Some morpho-species cannot be distinguished on the basis of DNA barcoding data alone. They might point to interesting biological cases of rapid reproductive isolation in sympatry or introgression. However, teasing out among these hypotheses requires additional data to sample a higher proportion of the genome.

These exceptions make identification from COI sequences alone challenging. They are however relatively uncommon, and barcoding will be accurate in species identification for most species with a success rate close of 90%. When morphological information is available, accuracy will be higher.

Our comprehensive dataset allowed us to explore geographic patterns of diversification. Divergence in allopatry is the most common mode of speciation, but sea cucumbers show high proportion of closely related species occurring in sympatry, a pattern that differs from other marine invertebrates. The mechanisms underlying this pattern are still unknown but rapid evolution in gamete recognition proteins seems to be driving similar patterns in other groups of echinoderms, but remain to be identified in sea cucumbers.

## 6 Supplementary material

**Figure 9:**
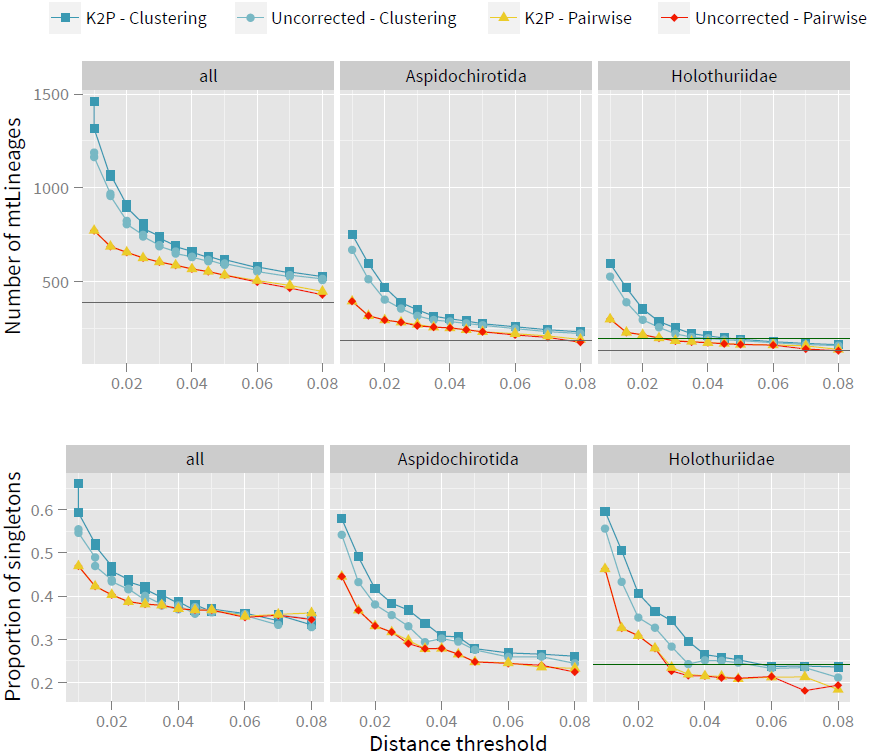
Comparison of the number of mtLineages (top), and the proportion of mtLineages represented by a single individual (singletons, bottom) estimated with the pairwise distance method (pairwise) and the clustering method (cluster), based on uncorrected distances (uncorrected) orthe Kimura 2-parameter distances (K2P), for the entire dataset (all), the order Aspidochirotida and the family Holothuriidae. The horizontal dashed line represents the numberof named species sequenced, the dotted line represents the number of mtLineages (top), and the proportion of singleton (bottom) delineated manually.

**Figure 10:**
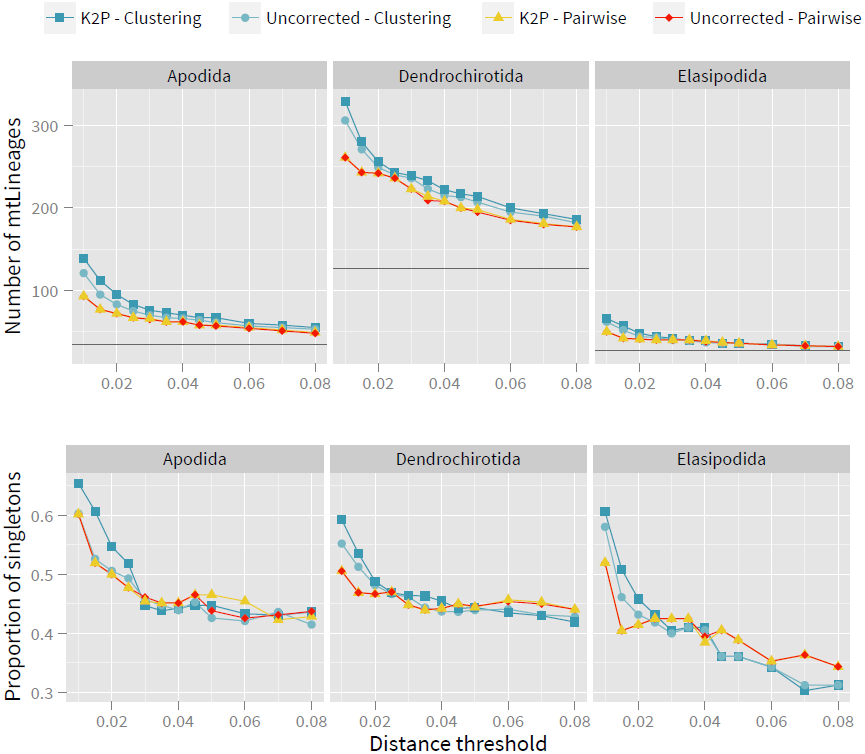
Comparison of the number of mtLineages (top), and the proportion of mtLineages represented by a single individual (singletons, bottom) estimated with the pairwise distance method (pairwise) and the clustering method (cluster), based on uncorrected distances (uncorrected) orthe Kimura 2-parameter distances (K2P), for the orders Apodida, Dendrochirotida and Elasipodida. The gray horizontal dashed line represents the number of named species sequenced.

